# Oral ingestion of the environmental toxicant trichloroethylene in rats induces alterations in the gut microbiome: relevance to idiopathic Parkinson’s disease

**DOI:** 10.1101/2022.02.19.481161

**Authors:** Neda M. Ilieva, Zachary D. Wallen, Briana R. De Miranda

## Abstract

Microbial alterations within the gut microbiome appear to be a common feature of individuals with Parkinson’s disease (PD), providing further evidence for the role of the gut-brain axis in PD development. As a major site of contact with the environment, questions have emerged surrounding the cause and effect of alterations to the gut microbiome by environmental contaminants associated with PD risk, such as pesticides, metals, and organic solvents. Recent data from our lab shows that ingestion of the industrial byproduct and environmental pollutant trichloroethylene (TCE) induces key Parkinsonian pathology within aged rats, including the degeneration of dopaminergic neurons, α-synuclein accumulation, neuroinflammation, and endolysosomal deficits. As TCE is the most common organic contaminant within drinking water, we postulated that ingestion of TCE associated with PD-related neurodegeneration may alter the gut microbiome to a similar extent as observed in persons with PD. To assess this, we collected fecal samples from adult rats treated with 200 mg/kg TCE over 6 weeks via oral gavage – the dose that produced nigrostriatal neurodegeneration – and analyzed the gut microbiome via whole genome shotgun sequencing. Our results showed changes in gut microorganisms reflective of the microbial signatures observed in individuals with idiopathic PD, such as decreased abundance of short-chain fatty acid producing *Blautia* and elevated lactic-acid producing *Bifidobacteria*, as well as genera who contain species previously reported as opportunistic pathogens such as *Clostridium*. From these experimental data, we postulate that TCE exposure within contaminated drinking water could induce alterations of the gut microbiome that contributes to chronic disease risk, including idiopathic PD.

## Introduction

Trichloroethylene (TCE) and structurally similar chlorinated solvents are pervasive environmental toxicants that contaminate air, water, and soil throughout the world (Bonvallot N, 2010; Cancer, 2014; Liu et al., 2020). Restrictions on TCE use within the United States and Europe have reduced its overall effluence into the environment in recent years, however the US Agency for Toxic Substances and Disease Registry (ATSDR) reports that TCE is the most frequently reported organic groundwater contaminant in the US, estimated to be present in 9 to 34% of drinking water (Todd et al., 2019). In addition, use of TCE as a degreasing agent in food production machinery results in contamination of certain processed foods, the most significant of which are butter and margarine, which averaged 73.6 *μ*g/kg when sampled in 1995 (USEPA, 2001). More recently, TCE was detected above 0.4 ppb in drinking water from 438 utilities over 43 states in the US from 2017-2019 (Environmental Working Group, 2022). Despite the reduction of TCE usage in some nations, production of TCE remains at a steady global growth, resulting in reports of heavily contaminated drinking water as recently as 2020 within certain global regions, such as the Lanzhou district of China (Liu *et al*., 2020). Because TCE is environmentally persistent, it will continue to be a global contaminant of concern for decades to come.

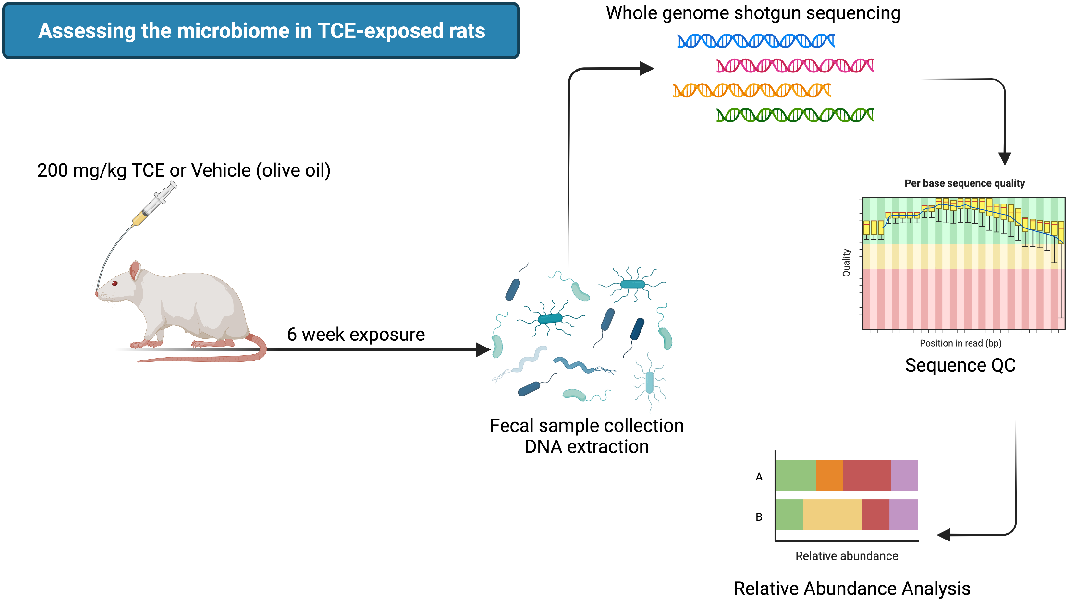

As risk for chlorinated solvent ingestion remains in place, questions about the effect on the gut microbiome and its role in chronic disease have arisen. TCE exposure is associated with autoimmune diseases, such as systemic lupus erythematosus and autoimmune hepatitis, which are linked to altered gut microbiome composition (Anagnostopoulos et al., 2004; Kamijima et al., 2007). Experimental models of autoimmune disease from TCE exposure show that ingestion of low levels of the solvent within drinking water (e.g., 0.5 mg/mL) resulted in gut microbiome alterations such as increased abundance of *Akkermansiaceae* and *Lachnospiraceae* families (Wang et al., 2021b). In addition, TCE-induced microbiome changes were associated with altered gut cytokine levels, elevated colonic oxidative stress, and decreased intestinal epithelial tight junction protein expression (Blossom et al., 2020; Khare et al., 2019; Wang *et al*., 2021b).

Epidemiological evidence also links chlorinated solvent exposure, such as TCE, tetrachloroethylene (PERC), and methylene chloride, with elevated risk for Parkinson’s disease (PD) (Goldman et al., 2012; Nielsen et al., 2021). Similarly, rodent studies of TCE exposure, including our own, recapitulate multiple features of PD pathogenesis, including the degeneration of nigrostriatal dopaminergic neurons (Liu et al., 2010; Liu et al., 2018), glial inflammation, endolysosomal deficits, and the accumulation of α-synuclein (De Miranda et al., 2021). As increasing evidence supports a role for the gut microbiome in the complex etiology of environmental, lifestyle, and genetic factors that contribute to idiopathic PD (Romano et al., 2021; Toh et al., 2022; Wallen et al., 2020), we postulated that TCE ingestion may induce gut microbiome alterations that influence neurodegeneration.

Previously reported data from our lab shows that aged rats exposed to 200 mg/kg TCE via daily oral gavage (or control, olive oil) for six weeks results in selective degeneration of nigrostriatal dopaminergic neurons, elevated nigral oxidative stress, endolysosomal impairment, and a-synuclein accumulation in the substantia nigra (De Miranda *et al*., 2021). To assess the effect of TCE on the gut microbiome at the same dose and schedule that induces Parkinsonian neurodegenerative pathology, we treated aged adult, female Lewis rats with 200 mg/kg TCE or vehicle (olive oil) via oral gavage for six weeks and collected fecal samples for whole genome shotgun sequencing. The resulting metagenomic analysis showed that TCE ingestion induced significant alterations in the gut microbiome. Some of the microbial signatures detected mirrored those observed in individuals with idiopathic PD, such as decreased abundance in the butyrate producing genus *Blautia*, and elevated abundance of lactic-acid producing *Bifidobacteria* and genera who contain species previously reported as opportunistic pathogens such as *Clostridium* (Romano *et al*., 2021; Wallen *et al*., 2020). In addition, some species-level alterations detected by metagenomic sequencing in TCE exposed rats were reflective of reported changes in genera from PD gut microbiota, such as *Parasutterella (Mao et al., 2021), Adlercreutzia* (Zhang et al., 2020), *and Mucispirillum* (Lin et al., 2019). Collectively, these data suggest that TCE exposure associated with nigrostriatal Parkinsonian pathology, also alters the gut microbiome in rats with similarities observed in idiopathic PD. As an environmental contaminant with relevant ingestion risk, we suspect that chronic alteration of the gut microbiome by TCE exposure may influence PD etiology via the gut-brain axis.

## Materials and Methods

### Reagents

Standard laboratory reagents and chemicals were purchased from Millipore Sigma (Burlington, MA) or Thermo Fisher Scientific (Waltham, MA). Trichloroethylene was purchased through Thermo Fisher Scientific (CAS #79-01-6), 99.6% purity. Premium olive oil was purchased at Trader Joes (Monrovia, CA).

### Animals and treatment

Female Lewis rats aged 11 months were purchased through a retired breeding program at Envigo Biosciences (Indianapolis, IN) and transported to the University of Alabama at Birmingham small animal facility for a one-week acclimation period prior to study onset. Female rats used in this study weighed an average of 251.33 g. Rats were provided with standard rodent chow and filtered water ad libitum throughout the study period, were double-housed with platforms and wooden blocks for enrichment, and maintained at 72-74°C with 50-60% humidity. Following a one-week handling period, rats were given a daily oral gavage of 200 mg/kg TCE or vehicle (olive oil) using flexible polypropylene gavage tubes (Instech, Plymouth Meeting, PA), 6-week period as reported in (De Miranda et al., 2021), as this dose and schedule produced the selective neurodegeneration of dopaminergic neurons from the nigrostriatal tract in both male and female aged Lewis rats. At the study endpoint, rats were placed inside of sterilized microisolator cages where fecal matter was collected and immediately flash frozen and stored at −80°C. At the study termination, rats were humanely euthanized using a lethal dose of pentobarbital euthanasia, and underwent transcardial perfusion with PBS, followed by organ collection. All animal experiments were conducted with approval by the University of Alabama at Birmingham Institutional Animal Care and Use Committee (IACUC).

### DNA extraction, sequencing, and bioinformatics

Frozen fecal samples were sent to CosmosID (Germantown, MD) for DNA extraction and whole genome (metagenome) shallow shotgun sequencing. DNA from samples was isolated using the QIAGEN DNeasy PowerSoil Pro Kit, according to the manufacturer’s protocol. Samples were quantified using the GloMax Plate Reader System (Promega) using the QuantiFluor® dsDNA System (Promega). DNA libraries were prepared using the Nextera XT DNA Library Preparation Kit (Illumina) and IDT Unique Dual Indexes with total DNA input of 1ng. Genomic DNA was fragmented using a proportional amount of Illumina Nextera XT fragmentation enzyme. Unique dual indexes were added to each sample followed by 12 cycles of PCR to construct libraries. DNA libraries were purified using AMpure magnetic Beads (Beckman Coulter) and eluted in QIAGEN EB buffer. Libraries were then sequenced on an Illumina HiSeq X platform 2×150bp.

Unassembled sequencing reads were directly analyzed by CosmosID bioinformatics platform for multi-kingdom microbiome analysis and profiling of antibiotic resistance and virulence genes and quantification of organisms’ relative abundance as described in (Hasan et al., 2014; Lax et al., 2014; Ottesen et al., 2016; Ponnusamy et al., 2016). Briefly, the system utilizes curated genome databases and a high-performance data-mining algorithm that rapidly disambiguates hundreds of millions of metagenomic sequence reads into the discrete microorganisms engendering the particular sequences. Similarly, the community resistome and virulome, the collection of antibiotic resistance and virulence associated genes in the microbiome, were also identified by querying the unassembled sequence reads against the CosmosID curated antibiotic resistance and virulence associated gene databases. Functional profiling and relative abundance from microbiome samples were identified using the strain-level curated CosmosID bacteria database, available through the CosmosID Bioinformatics Platform (CosmosID Metagenomics Cloud, app.cosmosid.com, CosmosID, Inc. www.cosmosid.com). Taxonomic profiles were filtered by the CosmosID Bioinformatics Platform using a filtering threshold that considers significant results based on internal statistical scores determined by analyzing a large number of diverse metagenomes. The resulting relative abundance data for detected bacteria was downloaded from the CosmosID platform and subsequently log transformed for futher analyses.

### Data Sharing

Raw metagenomic sequencing data from this study is deposited in Mendeley Data; De Miranda, Briana (2022), “Gut microbiome metagenomic data - oral ingestion of TCE in rats”, Mendeley Data, V1, doi: 10.17632/j3xy2wz3z2.1. Relative abundance averages and fold change calculations from all measured species in the gut from vehicle (VEH) and TCE treated rats are available log-transformed and untransformed data in Supplemental Material (Table S1).

### Data analysis

Statistical analyses were conducted using R Studio, GraphPad Prism 5.0 software (GraphPad Software Inc., San Diego, USA), and CosmosID Bioinformatics Platform as indicated. Principal components analysis (PCA), visualization of the PCA, and graphics were conducted using the ‘FactoMineR’ version 2.4 and ‘factoextra’ version 1.0.7 packages for R. PCA was conducted on log transformed relative abundances of bacterial species, using the PCA function from ‘FactoMineR’ and visualized via the ‘fviz_pca_ind’ function from ‘factoextra’. PCA grouping was based on the treatment group (VEH or TCE) and ellipses were created for the confidence intervals (95% CI) of each group. Permutational multivariable analysis of variance (PERMANOVA) was conducted using the ‘vegan’ package in R version 2.5-7 via the adonis function.

Shannon and Simpson indices for alpha-diversity were calculated using the ‘diverse’ package version 1.9.90 function ‘diversity’ in R for both VEH and TCE groups. Data was exported then plotted in GraphPad Prism Software. Statistical differences between log transformed relative abundance of bacterial genera from VEH and TCE animals were determined using t-test correcting for multiple comparisons by controlling the false discovery rate (Benjamini and Hochberg) using Graphpad Prism. Linear discriminant analysis (LDA) effect size (LEfSe) was used to ascertain taxonomic differences in untransformed relative abundances of bacterial species from VEH and TCE groups, performed with the CosmosID Bioinformatics Platform. Only taxa with p < 0.05 (Kruskal-Wallis test) were compared for effect size using LDA, and features that reached LDA > 2 were plotted as discriminatory features between treatment groups. Comparisons of relative abundance within bacterial genera and species from VEH or TCE treated rats were conducted using an unpaired student’s t-test using Graphpad Prism.

## Results

### TCE ingestion influences gut microbiome composition

Two-dimensional principal component analysis (PCA) based on log-transformed relative abundances of bacterial species revealed a separation between VEH and TCE treated rats. Differences in gut microbiome compositions observed in PCA were statistically significant (*p* = 0.008), tested using PERMANOVA (Figure 1a). Comparable species alpha-diversity was observed in TCE and VEH groups measured with Simpson and Shannon indices (**Figure 1b-c**). Bacterial phyla of VEH and TCE animal gut microbiota with 1% or more relative abundance are depicted in Figure 2a, with no significant differences being detected between taxa (*p* = 0.77). At the genus level, TCE ingestion caused a reduction *Ligilactobacillus* (*p* < 0.01) compared to VEH (**Figure 2b-c**). Using the linear discriminant analysis (LDA) effect size (LEfSe) analysis, we identified differential relative abundance in several features between VEH and TCE treated animals that were considered statistically significant with *p* < 0.05 (Kruskal-Wallis test) and LDA score > 2 (**Figure 3**).

**Figure 1.**
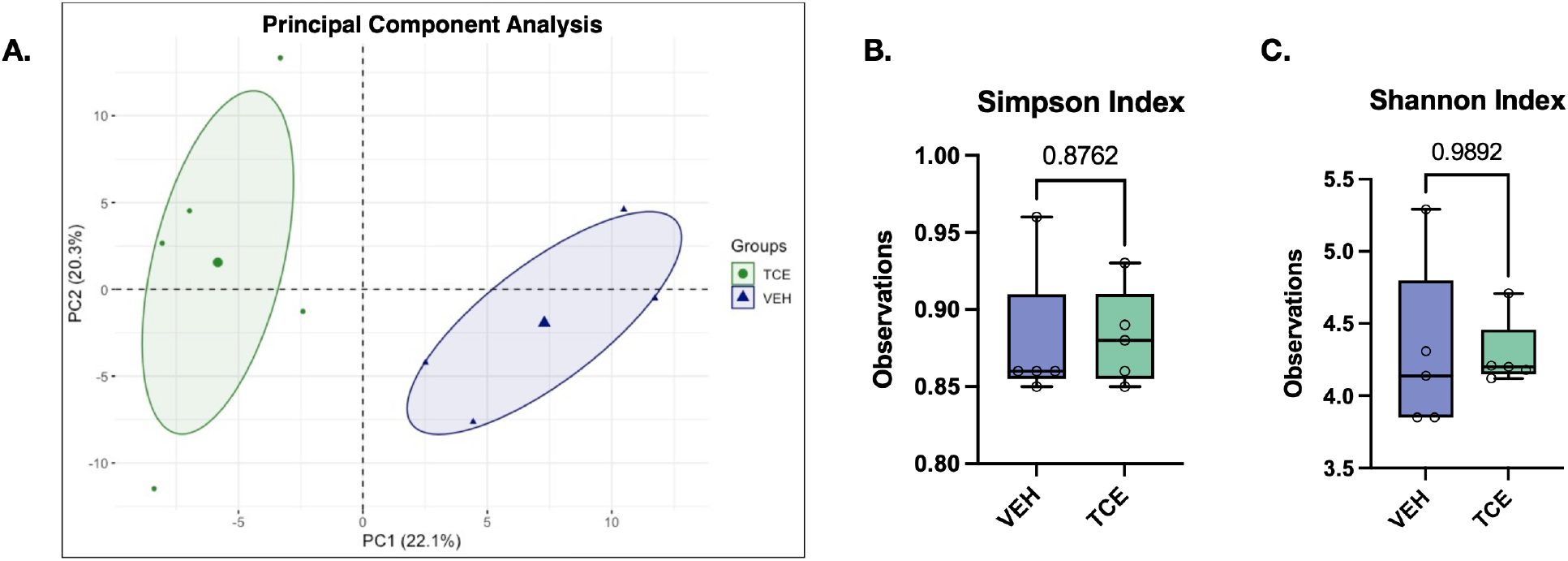
Gut microbiome composition measured by relative abundance differs between animals exposed to TCE and VEH. **A**. 2-Dimensional Principal Component Analysis (PCA) of relative abundances for all bacterial species sequenced, shows separation and independent clustering of TCE (green) and VEH (blue) groups, p = 0.008 (PERMANOVA). **B-C.** Simpson and Shannon indices demonstrate comparable diversity between microbiota from TCE and VEH animals. N=5, box and whisker lines denote median and interquartile range.

**Figure 2.**
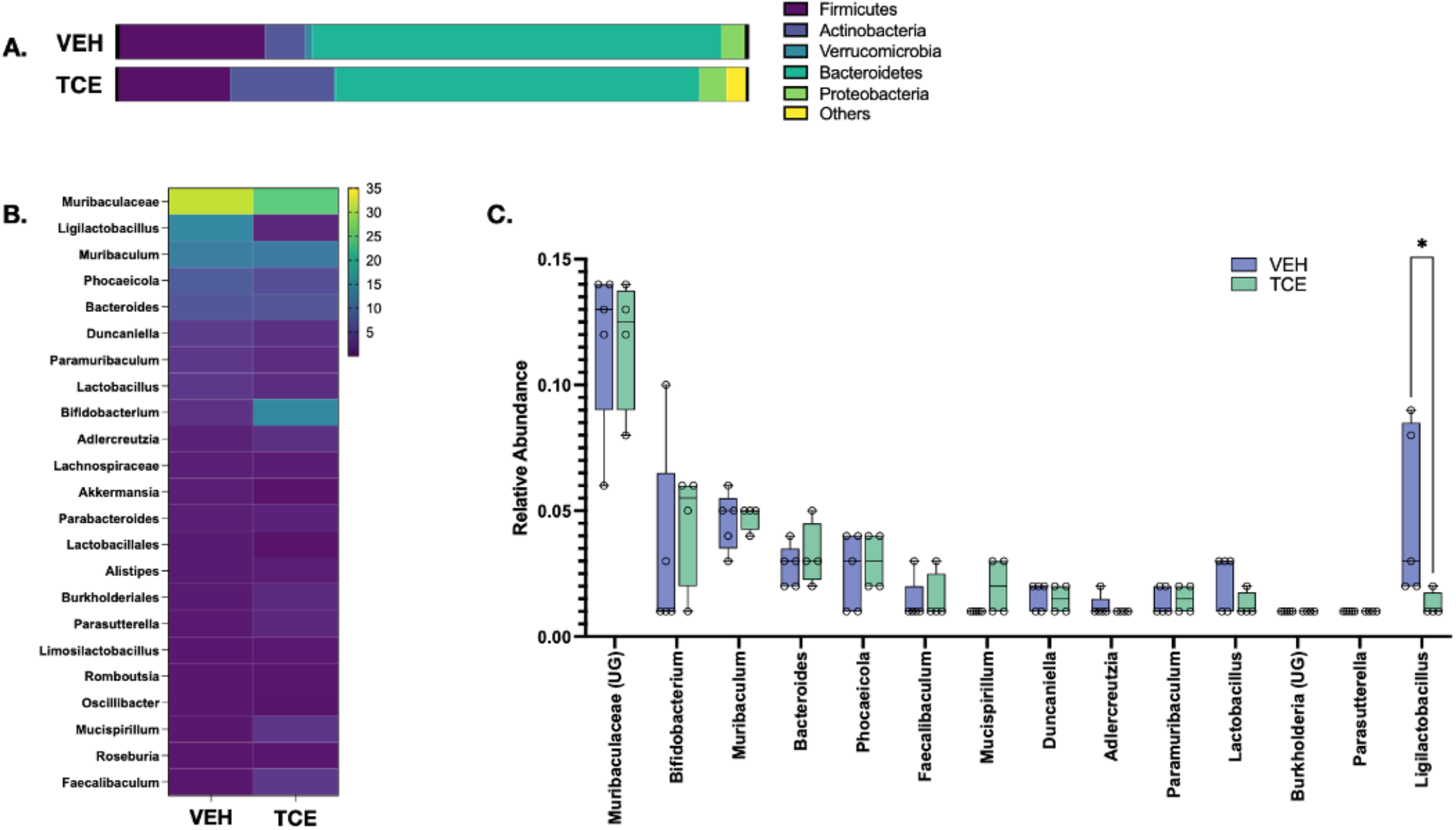
Relative abundance of gut microbiome genera is influenced by TCE exposure in rats. **A**. Graphical representation of log-tranformed relative abundances of phyla in VEH and TCE groups, with no statistical differences between taxa. **B.** Heatmap depiction of log-transformed relative abundance of major gut microorganisms from VEH and TCE treated rats, scale bar indicates % relative abundance. **C.** A comparison between log-transformed relative abundance of bacterial genera from the microbiome of VEH and TCE rats with significantly decreased *Ligilactobacillus* (p = 0.001); N= 5, multiple comparison t-tests with FDR, \*\**p* < 0.01, box and whisker lines denote median and interquartile range.

**Figure 3.**
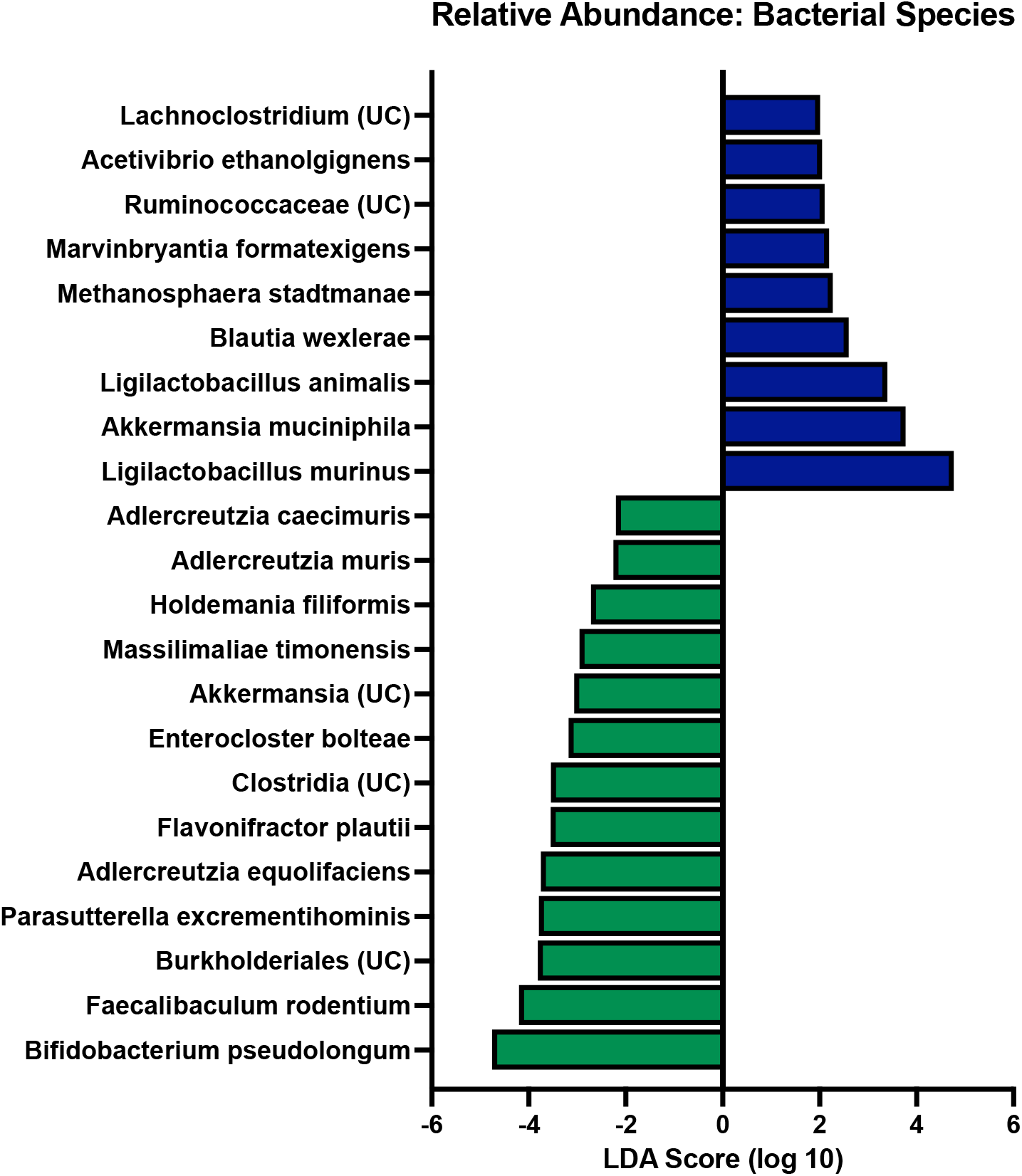
TCE ingestion resulted in significant alteration of gut microbiome bacterial species relative abundance. Linear discriminant analysis effect size (LEfSe) demonstrates relative abundance differences between metagenomic sequences of bacterial species from VEH (blue) and TCE (green) treated rats. Unclassified species of a bacterial genus are denoted as (UC).*p* < 0.05 LDA score >2, N =5.

### Microbial signatures in TCE treated rats share similarities with some idiopathic PD gut microbiome patterns

We identified several trends in bacterial species from TCE treated rats that matched published reports of microbiome composition reported in individuals with idiopathic PD compared to healthy controls. A decrease in the short-chain fatty acid (SCFA) producing bacteria *Blautia* was a prominent feature of TCE treated rat microbiome compared to VEH (*p* < 0.05), which is consistently reported in idiopathic PD gut microbiota (Romano *et al*., 2021; Wallen *et al*., 2020), as well as other chronic diseases (Liu et al., 2021; Ren et al., 2020) (**Figure 4a**). Bacterial genera such as *Clostridium* (*p* < 0.05) and *Bifidobacterium* (*p* < 0.05), which are observed in greater abundance in the gut microbiota of PD patients (Gerhardt and Mohajeri, 2018a; Hill-Burns et al., 2017b; Romano *et al*., 2021; Wallen *et al*., 2020), were significantly increased in TCE treated animals compared to VEH (**Figure 4b-d**). A type of unclassified *Burkholderiales* species which belong to *Burkholderia*, a bacterial genus involved in generating hydrophobic bile acids (Wallner et al., 2019), displayed elevated abundance in TCE treated rats (*p* < 0.001), sharing a similar pattern with observations made from genera in the PD appendix (Li et al., 2021). Bacterial species *Parasutterella excrementhominis* (*p* < 0.001), *Adlercreutzia equolifaciens* (*p* < 0.05), and *Muscipirillum schaedleri* (*p* < 0.05), and *Holdemania filiformis* (*p* < 0.01) were also observed as significantly elevated in TCE treated rats which reflects what has been observed at the genus level from the PD gut microbiome (Lin *et al*., 2019; Mao *et al*., 2021; Zhang *et al*., 2020) (**Figure 4e-h**). A comparison of log-transformed differentially abundant microbiota in TCE exposed rats (versus VEH) with reported alterations in PD (versus health control) is described in **Table 1.**

**Figure 4.**
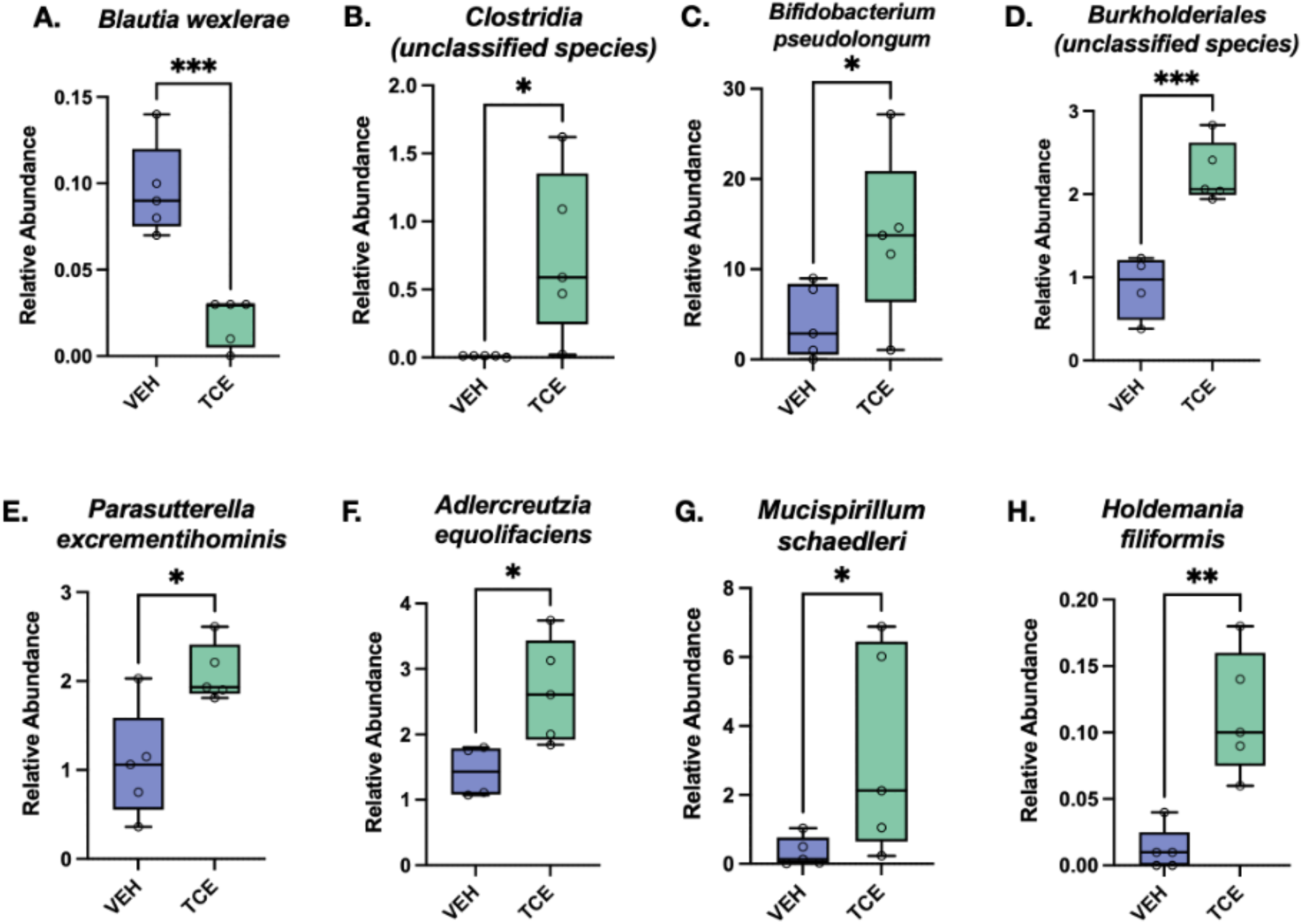
Relative abundance of gut microbial species within TCE-treated rats share similarities with alterations reported in PD. TCE treatment resulted in reduction of *Blautia* (*p* = 0.02; **A**), and greater relative abundance of an unspecified species of *Clostridia* (*p* = 0.025), *Bifidobacterium* (*p* = 0.031), and *Burkholderiales* species (*p* = 0.001), *Parasutterella excrementihominis* (*p* = 0.009), *Adlercreutzia equolifaciens* (*p* = 0.025), *Mucispirillum schaedleri* (*p* = 0.031), and *Holdemania filiformis* (*p* = 0.0017) compared to VEH animals, which are reported trends in PD gut microbiota (**B-H**). N=5, \**p* < 0.05, \*\**p* < 0.01, \*\*\**p* < 0.001, box and whisker lines denote median and interquartile range.

**Table 1.** Differentially abundant bacteria from TCE treated rats in relation to microbiota in persons with PD. Species data are reported from log-transformed metagenomic analyses in TCE treated rats belonging to genera in 16S rRNA or targeted gene sequencing from human data. Unclassified species belonging to a genera are labed as (uc).

| Bacterial Species | TCE Microbiome | Abundance in Parkinson's Patients (Associated Genus or Species) |  |
| --- | --- | --- | --- |
| | Log Fold $\Delta$, $p$ value | Increased | Reduced |
| *Blautia wexlerae* | -0.345, $p = 0.0472$ | ↑ (Heintz-Buschart *et al.*, 2018; Li *et al.*, 2019; Scheperjans *et al.*, 2015) | ↓ (Aho *et al.*, 2019; Barichella *et al.*, 2019; Hill-Burns *et al.*, 2017b; Petrov *et al.*, 2017; Takahashi *et al.*, 2022; Zapala *et al.*, 2021) |
| *Clostridium (uc)* | 0.210, $p = 0.0513$ | ↑ (Heintz-Buschart *et al.*, 2018; Kang *et al.*, 2020; Keshavarzian *et al.*, 2015; Lin *et al.*, 2018; Lubomski *et al.*, 2020; Murros, 2021; Murros *et al.*, 2021; Qian *et al.*, 2018; Romano *et al.*, 2021) | ↓ (Aho *et al.*, 2019; Bedarf *et al.*, 2017; Hasegawa *et al.*, 2015) |
| *Bifidobacterium pseudolongum* | 0.682, $p = 0.064$ | ↑ (Barichella *et al.*, 2019; Heintz-Buschart *et al.*, 2018; Hill-Burns *et al.*, 2017a; Lin *et al.*, 2018; Petrov *et al.*, 2017; Romano *et al.*, 2021; Unger *et al.*, 2016) | |
| *Burkholderiales (uc)* | 0.210, $p = 0.0280$ | ↑ (Li *et al.*, 2020a) | |
| *Parasutterella excrementihominis* | 0.205, $p = 0.0280$ | ↑ (Heintz-Buschart *et al.*, 2018; Heinzel *et al.*, 2021; Mao *et al.*, 2021; Tan *et al.*, 2018) | ↓ (Hill-Burns *et al.*, 2017a) |
| *Adlercreutzia eqolifaciens* | 0.143, $p = 0.0165$ | ↑ (Jin *et al.*, 2019; Zhang *et al.*, 2020) | |
| *Mucispirillum schaedleri* | 0.359, $p = 0.0750$ | ↑ (Lin *et al.*, 2019; Rajput *et al.*, 2021) | |
| *Akkermansia muciniphila* | -1.077, $p = 0.0086$ | ↑ (Barichella *et al.*, 2019; Bedarf *et al.*, 2017; Heintz-Buschart *et al.*, 2018; Hill-Burns *et al.*, 2017a; Keshavarzian *et al.*, 2015; Li *et al.*, 2019; Romano *et al.*, 2021; Unger *et al.*, 2016) | ↓ (Aho *et al.*, 2019) |
| *Ligilactobacillus animalis/murinus* | -0.805, -0.938; $p = 0.0009$, 0.011 | ↑ (Aizawa *et al.*, 2019; Romano *et al.*, 2021) | |
| *Holdemania filiformis* | 0.352, $p = 0.0250$ | ↑ (Mao *et al.*, 2021; Qian *et al.*, 2018) | |

### Functional profiling of biochemical activities from relative abundance

Whole genome shotgun sequencing data was analyzed for functional biochemical activities using the MetaCyc Pathways database via the CosmosID Bioinformatics Platform. TCE exposed rats displayed significantly increased activity in pathways involving ATP synthesis and hydrolysis (*p* = 0.0086, *p* = 0.0076 compared to VEH, respectively). Though limited data are available to compare the effect of TCE with human PD microbiome functional pathways, some reports of similarly altered biochemical processes in TCE treated rats, such as ubiquinone biosynthetic processes, were observed in idiopathic PD and multiple systems atrophy (Engen et al., 2017; Keshavarzian *et al*., 2015) (**Figure 5**).

**Figure 5.**
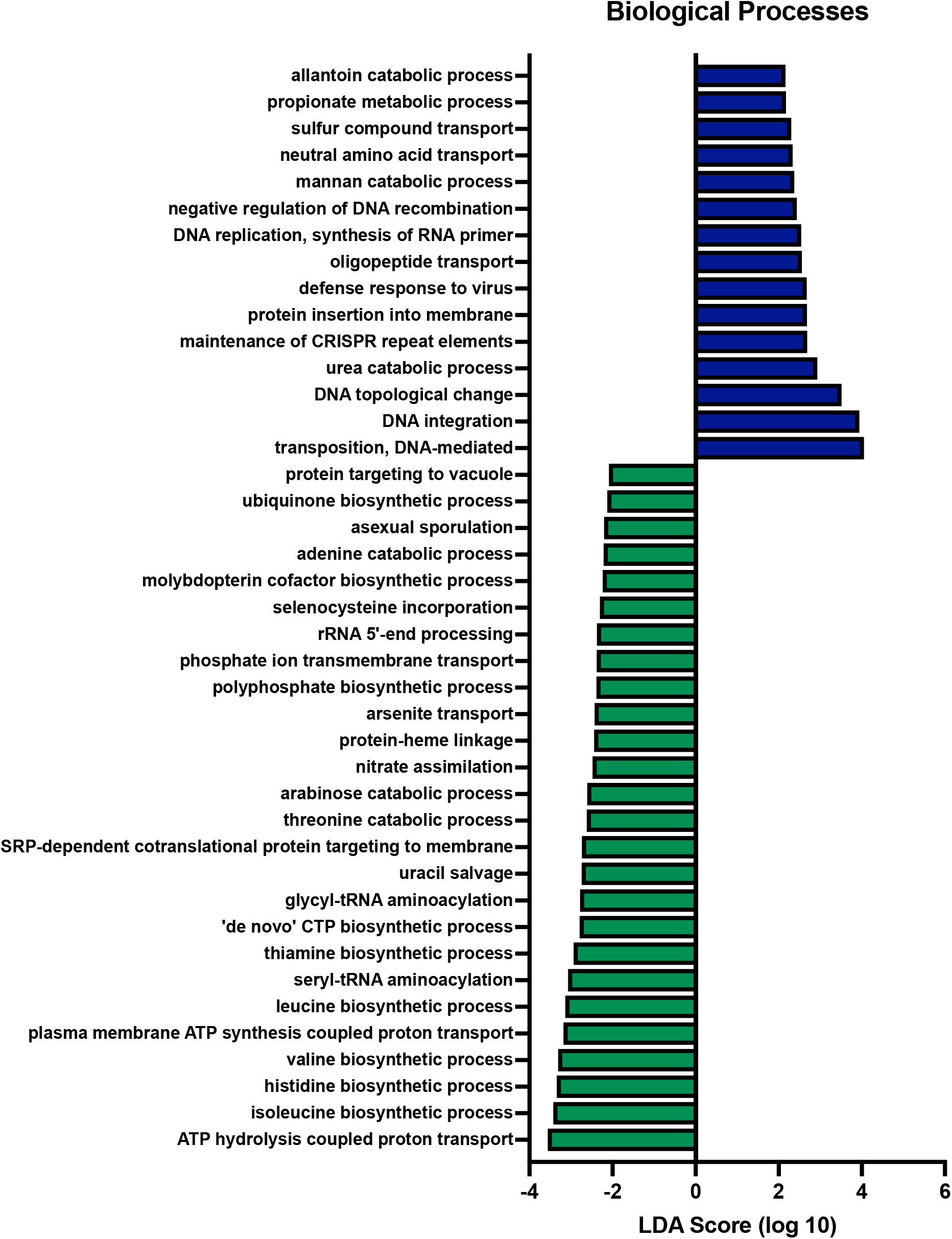
Predicted biological processes associated with TCE induced gut microbiome dysbiosis. LEfSe plot from pathway analysis data of biological processes (MetaCyC) shows significant differences in microbial biological processes between TCE and VEH rats. *p* < 0.05 LDA score > 2, N =5.

## Discussion

Exposure to environmental toxicants via ingestion may induce gut microbiome alterations that contribute to increased risk for adverse health outcomes and chronic disease (Popli et al., 2022). Despite the variability in human population study data (Keshavarzian et al., 2020), a gut microbiome signature appears to be present in most individuals with PD that includes the depletion in relative abundance of SCFA producing genera, such as *Blautia*, and increased relative abundance of *Bifidobacteria, Akkermansia, Clostridium*, and *Lactobacillus* (Heinzel *et al*., 2021; Hill-Burns *et al*., 2017b; Toh *et al*., 2022; Wallen *et al*., 2020). Thus, a convincing explaination suggests that altered gut microbiota contribute to elevated GI and systemic inflammation that may drive neuroinflammatory changes and/or influence the aggregation and spread of α-synuclein along the gut-brain axis (Houser and Tansey, 2017). In addition, the observed gut microbiome alterations in PD also indicate a role for environmental exposures that increase risk for the disease, such as pesticides, metals, air pollution, organic solvents, and pathogens, as the GI tract represents the most profuse site of environmental contact within the body (Keshavarzian *et al*., 2020).

At a dose shown to produce dopaminergic neurodegeneration from the nigrostriatal tract, exposure to 200 mg/kg TCE over 6 weeks in adult rats resulted in a profound dysbiosis in the gut microbiome, with a high degree of concordance to reported microbiota from idiopathic PD. The strong reduction in abundance of the SCFA producing genus *Blautia* from TCE-exposed rats is a similar feature to PD, as well as other chronic diseases, such as inflammatory bowel disease and ulcerative colitis, both of which are GI disorders associated with PD development (Villumsen et al., 2019). Conversely, significantly increased abundance of *Bifidobacterium, Clostridium*, and *Burkholderiales* was observed in TCE treated rats, which have been reported in one or more metagenomic or 16S datasets comparing the gut microbiome of healthy controls to individuals with PD (Keshavarzian *et al*., 2020; Keshavarzian *et al*., 2015; Li *et al*., 2021; Wallen *et al*., 2020). These data imply that similarities in the PD gut microbiome can be at least partially reproduced by exposure to a single environmental toxicant associated with elevated risk for the disease.

TCE is a highly lipophilic small molecule, which readily crosses biological membranes and is extensively metabolized by the liver and kidneys (Cichocki et al., 2016; Lash et al., 2006). Absorption of TCE or similar chlorinated solvents within the GI tract therefore may directly damage microorganisms and/or intestinal epithelial cells and proteins, resulting in elevated systemic inflammation that may ultimately influence PD development (Houser and Tansey, 2017; Khan et al., 2009). Epidemiological evidence exists for GI damage from chlorinated solvent exposure with structural similarities; for example, individuals who were exposed to public drinking water contaminated with tetrachloroethylene (PERC) had higher risk for colorectal cancer (Paulu et al., 1999). In addition, the association of TCE with autoimmune disease (Phillips et al., 2019) points to an intersection of GI pathology and microbiome dysbiosis from ingestion exposure that could influence inflammatory bowel diseases. Animal studies in the autoimmune mouse line MRL^+/+^ showed that exposure to TCE via drinking water resulted in irreversible gut microbiome dysbiosis, even after exposure cessation, specifically elevated *Bifidobacterium*, the genus found to be elevated to the greatest extent in this study (Khare *et al*., 2019). A separate study of TCE exposure in MRL^+/+^ mice showed TCE ingestion resulted in decreased expression of tight junction proteins ZO-2, occludin, and claudin-3, as well as elevated proinflammatory proteins CD14 and IL-1β in the colon (Wang *et al*., 2021b). Together, these data suggest that TCE ingestion induces a proinflammatory environment within the gut, with a possible bi-directional interplay between gut microbes, immune regulation, and intestinal pathology.

Though gut microbiome alteration is linked with TCE ingestion in animal models, it remains unclear what the functional implications of bacterial abundance mean in the context of environmental risk for PD. In the environment, TCE is metabolized by soil microorganisms, predominantly through reductive dichlorination by anaerobic bacteria (Freedman and Gossett, 1989). In soil contaminated by TCE, microbiota exhibit inhibited energy metabolism and a reduction in the metabolic capacity for other xenobiotics (Li et al., 2020b; Moccia et al., 2017). Thus, in addition to inflammatory promoting factors in the gut, TCE exposure could influence xenobiotic metabolism by gut microbiota, potentially resulting in altered enterohepatic cycling, host cytochrome P450 metabolizing enzyme expression, and reduced detoxification or elevated bioactivation. Interestingly, gut microbiota from TCE exposed rats displayed significantly increased activity in pathways involving ATP synthesis and hydrolysis, possibly related to elevated P450 metabolism (Foti and Dalvie, 2016). Gut microbiota from individuals with PD displayed elevated xenobiotic degradation pathways related to environmental contaminants such as pesticides and chloroalkanes (Hill-Burns *et al*., 2017b). In concordance with this, we observed elevated abundance of *Burkholderia* in TCE treated rats, a genus which contains species that specifically degrade polychlorinated biphenyls (Tehrani et al., 2012), and were reported to be increased in the PD appendix (Li *et al*., 2021). Collectively, these data support a role for environmental exposures potentially driving gut microbiome alterations observed in PD.

In line with the many other published studies investigating gut microbiome and PD related neurodegeneration, the data collected from TCE treated rats partially conflicted with human PD microbiome and other animal model studies (Keshavarzian *et al*., 2020). One of the major differences observed was the decrease in *Akkermansia* and *Lactobacillus* genera from TCE treated rats, compared to the relatively common increase found in human PD studies (Hill-Burns *et al*., 2017b; Romano *et al*., 2021; Wallen *et al*., 2020; Zapała *et al*., 2021), though at least one or more studies report decreases in either genus (Li *et al*., 2019; Qian *et al*., 2018). While the most obvious source of variation is likely rodent to human translation, MRL^+/+^ mice exposed to TCE in drinking water displayed a significant elevation in *Akkermansia* within the gut microbiome compared to control (Wang *et al*., 2021b). In addition, TCE exposed MRL^+/+^ mice displayed an apparent dose-dependent response in *Lactobacillus* abundance within the gut microbiome, as low dose TCE exposure resulted in increased *Lactobacillus* expression, however high doses of TCE were associated with its decrease after 259 days of exposure (Khare *et al*., 2019). As 200 mg/kg TCE in adult rats was a relatively high amount of TCE, reduced *Lactobacillus* abundance may be a result of dose-specificity. In relation, as the autoimmune animal studies show, elevated *Akkermansia* abundance induced by TCE exposure could be a result of host-microbiome-environment interaction involving autoimmunity (Khare *et al*., 2019; Wang et al., 2021a).

In addition to the disparate findings in certain genera, there were other limitations of this study. First, direct comparison of rats exposed to TCE and human PD is difficult; the inbred Lewis rats exposed in this study were fed a standard rodent chow and only underwent treatment with a single toxicant exposure over 6 weeks. In contrast, the microbiome from individuals with PD reported in **Table 1** is influenced by diet, medications, location of residence, and simultaneous or previous exposure to multiple environmental factors, which likely contributes to inconsistencies in the published literature which we used as a comparison (Gerhardt and Mohajeri, 2018b; Hill-Burns *et al*., 2017b; Keshavarzian *et al*., 2020; Romano *et al*., 2021). Variability between gut microbiome abundances in humans are also common, which are apparent from the differences in increased and decreased abundane of certain genera reported in publications listed in **Table 1**. Beyond diet, temporal differences have been observed in gut microbial abundance, with individual variability fluctuating significantly on a day-to-day basis, and up to 100-fold over a few weeks (Vandeputte et al., 2021). In addition, complex gene-environment interactions that occur in individuals with PD influence the gut microbiome (Wallen et al., 2021), which were not possible to measure in wildtype rats.

Due to sample size limitations to obtain metagenomic-scale microbiome data, only female rats were used for this study, therefore we were unable to measure sex differences, which may play a role in PD gut-brain interaction (Baldini et al., 2020; Cox et al., 2019). Finally, many comparisons were made between 16S sequencing reported in the literature and whole genome shallow shotgun sequencing performed here, possibly resulting in differences between curated microbiome databases and taxonomic resolution. Differences in data handling between human and animal microbiome studies also limited the direct comparison from our study to all others, however, we showed both log-transformed and untransformed data to make comparisons where appropriate. The magnitude of specific increased/decreased gut microbiota changes in TCE treated rats to humans cannot be assessed across each reported study in **Table 1** as not all studies reported log-transformed fold change values for each genus or species, however, we can infer some of overall extent from Wallen et al., 2020. As an example, *Blautia*, which we found to be reduced by 0.345 log fold change in TCE treated rats (*p* = 0.0472), was reduced in humans with PD by 0.68 log fold change (*p* = 2^−04^). Similarly, *Claustridia* was reported to be increased by 0.77 log fold change (*p* = 0.02) in human PD (Wallen *et al*., 2020), and 0.21 log fold change (*p* = 0.0513) in TCE treated rats.

Perhaps because of these limitations, the observed similarities in gut microbiota of rats exposed to TCE with reports from individuals with PD is quite remarkable and indicates a connection between the gut microbiome and relevant environmental contaminants associated with PD. Future studies to assess the influence of the gut microbiome on the chronic neurotoxic properties of TCE and other environmental toxicants will be essential in understanding PD risk.

## Acknowledgments

This work was supported by research grants from the National Institutes of Environmental Health Sciences (R00ES029986, BRD), and the Parkinson Association of Alabama (BRD).

